# Mechanically Resolved Imaging of Bacteria using Expansion Microscopy

**DOI:** 10.1101/622654

**Authors:** Youngbin Lim, Margarita Khariton, Keara M. Lane, Anthony L. Shiver, Katharine M. Ng, Samuel R. Bray, Jian Qin, Kerwyn Casey Huang, Bo Wang

## Abstract

Imaging dense and diverse microbial communities has broad applications in basic microbiology and medicine, but remains a grand challenge due to the fact that many species adopt similar morphologies. While prior studies have relied on techniques involving spectral labeling, we have developed an expansion microscopy method (μExM) in which cells are physically expanded prior imaging and their expansion patterns depend on the structural and mechanical properties of their cell walls, which vary across species and conditions. We use this phenomenon as a quantitative and sensitive phenotypic imaging contrast orthogonal to spectral separation in order to resolve bacterial cells of different species or in distinct physiological states. Focusing on host-microbe interactions that are difficult to quantify through fluorescence alone, we demonstrate the ability of μExM to distinguish species within a dense community through *in vivo* imaging of a model gut microbiota, and to sensitively detect cell-envelope damage caused by antibiotics or previously unrecognized cell-to-cell phenotypic heterogeneity among pathogenic bacteria as they infect macrophages.

## Introduction

Imaging of heterogeneous bacterial populations has broad applications in understanding the complex microbiota that exist on and within our bodies, as well as complex host-microbial interfaces, yet remains a significant challenge due to the lack of suitable tools for distinguishing species and identifying altered physiological states [1–3]. Analyses to date have mostly relied on spectral separation using fluorescence in situ hybridization (FISH) with probes designed to target 16S RNA sequences specific to certain taxa [4], or genetically engineered microbes that express distinct fluorescent proteins [5]. However, these methods are generally insensitive to physiological changes in bacterial cells that are often modulated by host environments and believed to be critical for the growth and spatial organizations of microbes [6,7].

The bacterial cell wall is a macromolecule responsible for shape determination in virtually all bacteria. Although little is known about the molecular architecture of the cell wall in most non-model organisms, its dimensions can vary widely, with the wall typically thick (tens of nanometers [8,9]) in Gram-positive species and thin (∼2-4 nm) in Gram-negative species [10], and their rigidity can vary across ∼10-100 MPa for Young’s modulus [11,12]. The cell wall also has various biochemical compositions [13] and exhibit distinct spatial patterns of cross-linking density [14], molecular organization [15,16], thickness [8,9], and stiffness [11,12], all of which depend on species and cell physiology. Thus, cell wall mechanics can potentially provide a contrast that is orthogonal to spectral separation in distinguishing species and even cellular physiological states. However, while cell-wall structure and mechanics have been measured by electron microscopy and atomic force microcopy [8,10,17], these methods are low-throughput and incompatible with dense bacterial populations or *in vivo* applications.

To address this challenge, we develop a new method that extends the application of expansion microscopy (ExM) to bacteria, particularly in the contexts of multispecies communities and infection. ExM is a recently developed optical imaging method that was designed to provide superior resolution to traditional fluorescence imaging, but its applications to bacteria have so far been limited [18]. ExM relies on simple chemistry in which biomolecules are anchored to hydrogel networks, and then membranes are permeabilized by detergents while proteins are digested by proteases to allow physical expansion. The expanded sample can be imaged through conventional optical microscopy and digitally compressed to gain resolution. While prior studies have focused on achieving uniform expansion of samples [19–21], we posited that differences in the expandability of the cell wall could provide imaging contrast that reflects its molecular structural and mechanical properties.

In this study, we show that the peptidoglycan cell wall requires additional treatments to break down for expansion. We demonstrate, with both *in vitro* and *in vivo* applications, that the contrast in cell-wall expandability is sufficient to resolve different bacterial species within dense communities. In addition, this method is sensitive enough to detect cell-wall damages due to antibiotics or host defensive responses that are otherwise difficult to capture using traditional imaging methods. We anticipate this method to enable future research in three major areas: (1) super-resolution imaging of bacterial subcellular components [22] without the need for expensive optics or special fluorophores; (2) high-content imaging that integrates both spectral and mechanical contrasts to dissect complex microbial communities; and (3) *in vivo* phenotyping of cell-wall mechanics and integrity [4,23].

## Results

### Expansion provides quantitative imaging contrast to distinguish bacterial species

To test whether the cell wall indeed restricts expansion in ExM, we imaged a mixture of two common symbiotic bacteria isolated from the gut of the fruit fly *Drosophila melanogaster*, *Acetobacter tropicalis* (GFP-labeled) and *Lactobacillus plantarum* (mCherry-labeled) [24], using a standard ExM protocol (**Fig. 1A**, top) [25]. We quantified the expansion ratio of individual cells based on cell width, because width can be measured precisely regardless of cellular orientation in three dimensions. We found that *A. tropicalis* and *L. plantarum* cells only expanded ∼1.9- and ∼1.2-fold, respectively (**Fig. 1B**, top), both of which were smaller than the 4- fold expansion expected from previous characterizations of ExM [19,25].

**Figure 1:**
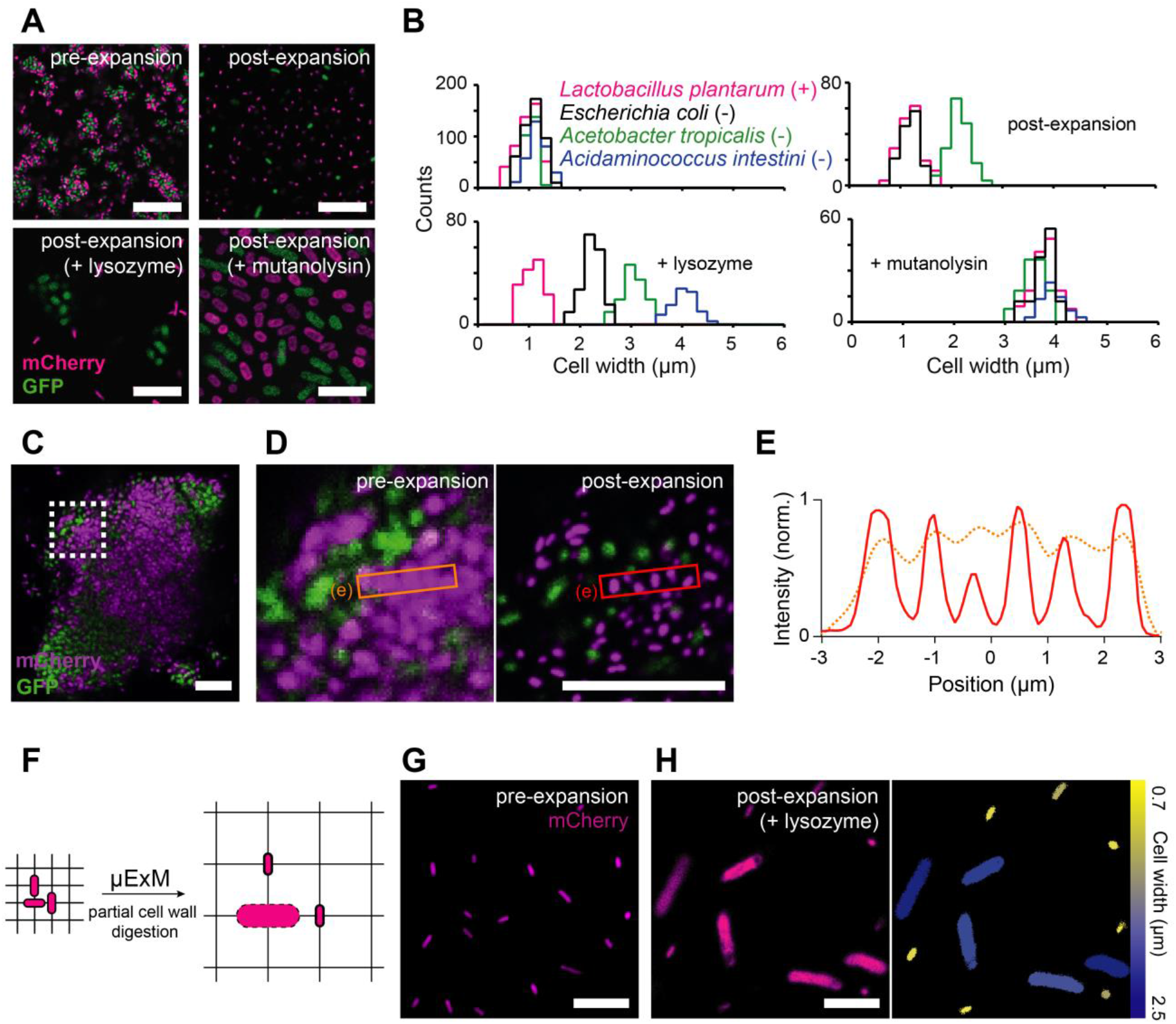
Differential expansion provides novel imaging contrast in μExM. (A) Representative confocal fluorescence images of mCherry-*L. plantarum* and GFP-*A. tropicalis* in close proximity. Top: untreated cells pre- and post-expansion; bottom left: cells treated with lysozyme to partially digest the bacterial cell wall before expansion; bottom right: cells treated with mutanolysin to fully digest the cell wall before expansion. (B) Quantification of cell width before and after expansion for representative microbial species. Lysozyme treatment maximizes the contrast in expansion between species, while mutanolysin treatment expands all species ∼4-fold. When fluorescently labeled strains were not available, we measured the expansion ratio using DNA staining. “(+)” and “(-)” denote Gram-positive and Gram-negative, respectively. (C) Pre-expansion image of a colony of mCherry-*L. plantarum* and GFP-*A. tropicalis*. (D) Magnified view of the boxed region in (C) before expansion (left) and after mutanolysin treatment and expansion (right). The scale bar in the post-expansion image is rescaled to match the pre-expansion dimension. (E) Cross-sectional normalized fluorescence intensity profiles of the boxed regions in (D), showing that μExM preserves the relative positions of cells (peaks in the orange and red curves overlap). (F) Schematic showing that differential expansion provides imaging contrast to distinguish microbial species in mixed populations. (G) Pre-expansion image of a mixture of mCherry-*L. plantarum* and mCherry-*E. coli*. The species have approximately the same dimensions, making them hard to distinguish. (H) μExM easily distinguishes *L. plantarum* and *E. coli* based on expansion ratio. Left: confocal image; right: the corresponding image false-colored by cell width. *E. coli* cells are expanded (blue) whereas *L. plantarum* cells remain unexpanded (yellow). All images are maximum intensity projections. Scale bars: 10 μm.

We then digested the wall using lysozyme from chicken egg white or mutanolysin from *Streptomyces globisporus*, after permeabilizing the cell membrane with methanol. Both enzymes are muramidases that cleave 1,4-beta-linkages between N-acetylmuramic acid and N-acetyl-D-glucosamine residues in the cell wall but differ in mechanism of action and activity [26]. Overnight lysozyme treatment enhanced the contrast in expansion ratios between species (**Fig. 1A,B**, bottom left): *A. tropicalis* cells were fully expanded ∼4-fold, whereas *L. plantarum* cells were largely unaffected. By contrast, mutanolysin treatment led to 4-fold expansion of both species (**Fig. 1A,B**, bottom right). The uniform expansion enabled us to resolve individual cells in densely packed mixed colonies of the two species while preserving their relative positions (**Fig. 1C–E**).

Next, we tested whether these findings could be extended to other commensal species from the human gut. We applied the method to seven species, including *Escherichia coli*, *Salmonella enterica*, *Acidaminococcus intestini*, *Bacteroides finegoldii*, *Parabacteroides distasonis*, *Bifidobacterium breve*, and *Clostridium innocuum*. These species were chosen to include both Gram-positives and Gram-negatives. For all species, the expansion ratios were approximately 4-fold after mutanolysin treatment, whereas the ratios after lysozyme treatment were generally larger for all Gram-negative species than for Gram-positives (**Table 1** and **S1 Fig.**), but varied widely, in particular among Gram-negatives (**Fig. 1B**, bottom left). Together, these results show that the extent of breakdown of the cell wall determines the expansion of bacterial cells, and the expansion ratio provides quantitative and fine resolution in distinguishing species beyond the tradition classification of Gram-negatives and Gram-positives. Henceforth, we refer to this method involving lysozyme or mutanolysin digestion as μExM, expansion microscopy of microbes.

**Table 1:** Expansion of bacteria is species specific.

| Strains<sup>1</sup> | Culture medium<sup>3</sup> | Culture condition | Antibiotic resistance | Expansion ratio<sup>4</sup> |
| --- | --- | --- | --- | --- |
| *Lactobacillus plantarum* | MRS | Aerobic, 30 °C | chloramphenicol | 1.12 ± 0.03 |
| *Acetobacter tropicalis* | MRS | Aerobic, 30 °C | tetracycline | 3.09 ± 0.07 |
| *Escherichia coli* | LB | Aerobic, 37 °C | ampicillin | 2.21 ± 0.04 |
| *E. coli imp4213* | LB | Aerobic, 37 °C | kanamycin | N.D. |
| *Salmonella enterica* (SL12023) | LB | Aerobic, 37 °C | ampicillin | 2.23 ± 0.05 |
| *Bifidobacterium breve*<sup>2</sup> | RCM | Anaerobic, 37 °C | N.A. | 1.21 ± 0.04 |
| *Clostridium innocuum*<sup>2</sup> | RCM | Anaerobic, 37 °C | N.A. | 1.14 ± 0.03 |
| *Acidaminococcus intestini*<sup>2</sup> | GAM | Anaerobic, 37 °C | N.A. | 3.98 ± 0.09 |
| *Bacteroides finegoldii*<sup>2</sup> | GAM | Anaerobic, 37 °C | N.A. | 3.67 ± 0.07 |
| *Parabacteroides distasonis*<sup>2</sup> | GAM | Anaerobic, 37 °C | N.A. | 3.79 ± 0.09 |

<sup>1</sup> magenta, Gram-positive; green, Gram-negative.
<sup>2</sup> obtained through BEI Resources, NIAID, NIH, as part of the Human Microbiome Project:
*Bifidobacterium breve* strain HPH0326, HM-856; *Clostridium innocuum* strain 6\_1\_30, HM-173; *Acidaminococcus intestini* strain D21, HM-81; *Bacteroides finegoldii* strain CL09T03C10, HM-727; *Parabacteroides distasonis* strain 31\_2, HM-169 (previously deposited as *Porphyromonas* sp.).
<sup>3</sup> MRS, De Man, Rogosa, and Sharpe; LB, Lysogeny Broth; RCM, Reinforced Clostridial Medium; GAM, Gifu Anaerobic Medium.
<sup>4</sup>The expansion ratio is computed as the average cell width in post-expansion images divided by the average cell width in pre-expansion images of lysozyme-treated cells, and reported as mean ± SEM (standard error of the mean).

Our findings suggest that μExM both improves imaging resolution, as shown in previous studies [19–21,25], and provides an additional imaging contrast associated with cell-wall mechanical properties that is orthogonal to spectral separation commonly used in fluorescence microscopy. We posited that in some cases, the contrast in expansion should be sufficient to distinguish between microbial species with differing cell-wall properties (**Fig. 1F**). Moreover, this contrast can be amplified by partial wall digestion using lysozyme. For example, the post-lysozyme expansion ratio distributions of four species, *L. plantarum*, *Escherichia coli*, *A. tropicalis*, and *Acidaminococcus intestini*, were clearly distinct, despite the variation across cells of the same genotype (**Fig. 1B**, bottom left). To directly test our hypothesis, we mixed populations of *L. plantarum* and *E. coli* cells, as they are of similar pre-expansion size and morphology and thus difficult to distinguish (**Fig. 1G**). Following lysozyme treatment and expansion, the two species could be easily separated by post-expansion size (**Fig. 1H**).

### Expansion detects cell-wall damage induced by antibiotics with high sensitivity

Given that expansion is dependent upon cell-wall properties, we reasoned that μExM should also reveal different expansion phenotypes in cells grown under conditions that generate cell-wall damage. Vancomycin is an antibiotic that binds to peptidoglycan precursors and prevents their crosslinking to the existing cell wall. While *E. coli* and most Gram-negative bacteria are typically resistant to vancomycin, the *imp4213* allele in the *lptD* gene, which disrupts synthesis of the lipopolysaccharide component of the outer membrane, leads to a permeable outer membrane and increased sensitivity to vancomycin [27]. Our previous results showed that vancomycin treatment of *imp4213* cells leads to the formation of pores in the cell wall from which the inner membrane and cytoplasm eventually escape when the pore size increases sufficiently [15]; before the point of this blebbing, it is difficult to detect the level of damage using existing light microscopy techniques.

We treated *imp4213* cells for 10 min with 1 μg mL^−1^ vancomycin; at this early time point, cells did not exhibit any morphological changes due to drug treatment. Unlike untreated controls that remained unexpanded (**Fig. 2A**), expansion of vancomycin-treated *imp4213* cells showed a striking pattern: surrounding the unexpanded cytoplasm (GFP-labeled) was a halo of DNA (TO-PRO-3-labeled) that occupied a space with width ∼4 times that of an unexpanded cell (**Fig. 2B,C**). We interpret this pattern as the translocation of DNA through pores in the cell wall during expansion (**Fig. 2D**). As the hydrogel network contained within the cell wall is unable to expand with the surrounding network, an extracellular cavity with low-density networks is created that lowers the effective chemical potential and drives the DNA to spread into this cavity (**Fig. 2E)**. We estimate the minimum chemical potential difference required for the spontaneous translocation of *E. coli* DNA to be ∼0.5 *k_B_T* (**Materials and Methods**), which is comparable to the thermal energy. This calculation indicates that the DNA translocation is a sensitive measure of cell-wall damage, as the DNA chain should always escape the confinement of the cell wall as long as the pore size grows to ∼100 nm (the Kuhn length of DNA). The size of these pores would be below the diffraction limit, and thus invisible using conventional optical approaches.

**Figure 2:**
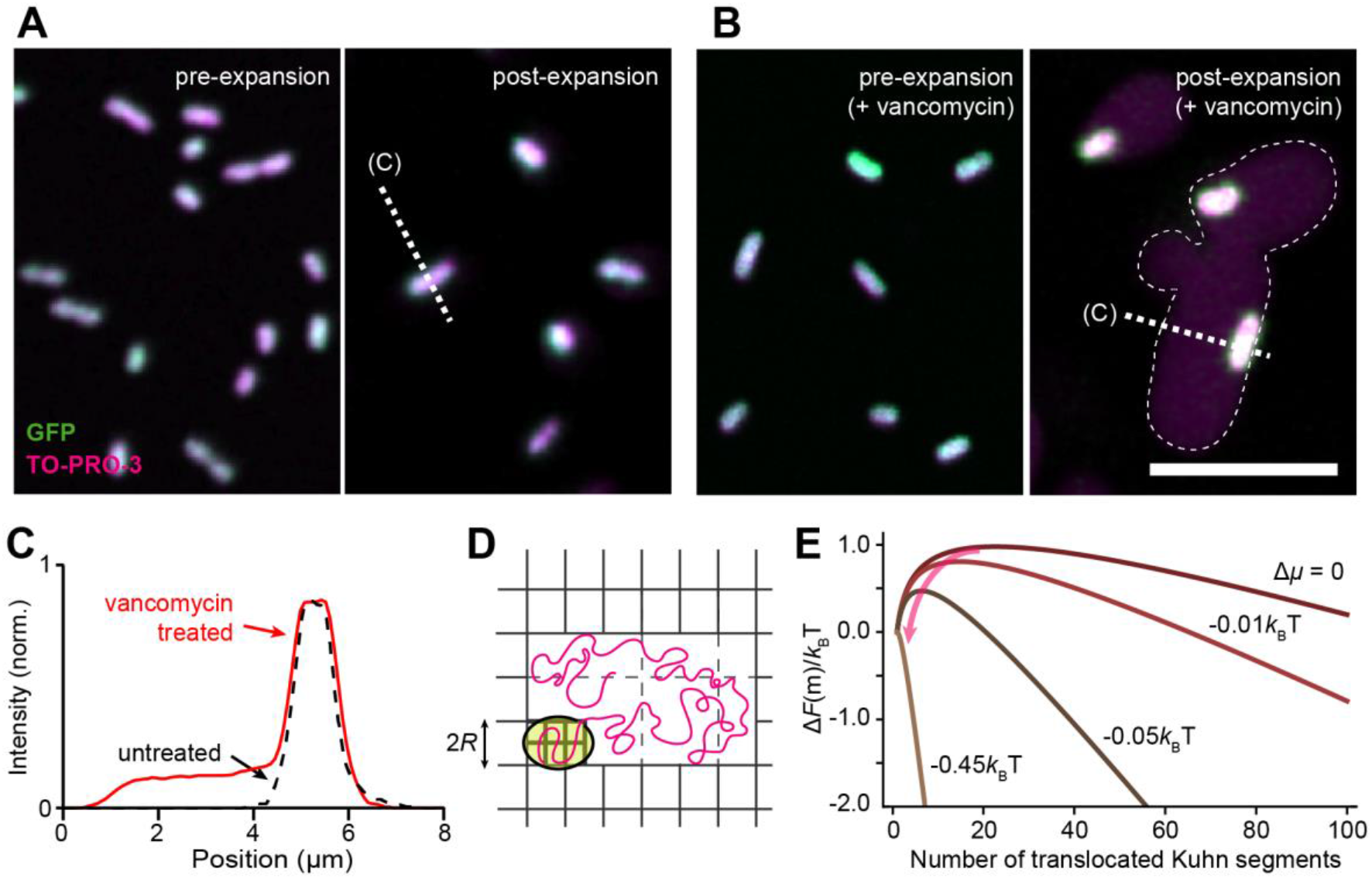
μExM detects with high sensitivity cell-wall damage induced by antibiotics. (A) Images of GFP-expressing *imp4213 E. coli* cells, with DNA co-stained using TO-PRO-3, before (left) and after (right) expansion. (B) Images of vancomycin-treated *imp4213* cells, in which expansion leads to a large halo of DNA fluorescence surrounding the unexpanded cytoplasm. All images are maximum intensity projections. Scale bars: 10 μm. (C) Normalized intensity profiles of TO-PRO-3 fluorescence measured along dashed lines in (A) and (B). (D) Proposed mechanism of DNA expansion through the translocation of a DNA chain. Yellow: cytoplasm; magenta: DNA; grey: gel network. Note the high-density gel network in the cell and low-density network around the cell. (E) The free energy for DNA translocation, Δ*F*(*m*), as a function of *m*^th^ Kuhn segment anchored at the pore. Δ*F*(*m*) has a maximum at *m*\*, which presents an entropic barrier for DNA translocation The lower density outside the cell wall due to expansion leads to a negative chemical potential difference (Δ*μ*), which facilitates the translocation process by reducing *m*\* (arrow) and eventually causes the entropic barrier to vanish (Δ*μ* < −0.45 *k*_B_*T*) for spontaneous DNA translocation.

### μExM resolves bacterial species within a model animal gut microbiota

To demonstrate the utility of expansion as an imaging contrast, we focused on two applications: (1) *in vivo* imaging to resolve bacterial species in an animal gut, particularly when strain-specific fluorescent tags may not be available or are limited by host tissue autofluorescence (**Fig. 3**), and (2) detection of cell wall disruption *in situ* when pathogenic bacteria are under attack from host defense mechanisms (**Fig. 4**). In both cases, traditional spectral contrast in fluorescence microscopy would be insufficient, and μExM reveals new biological insights.

**Figure 3:**
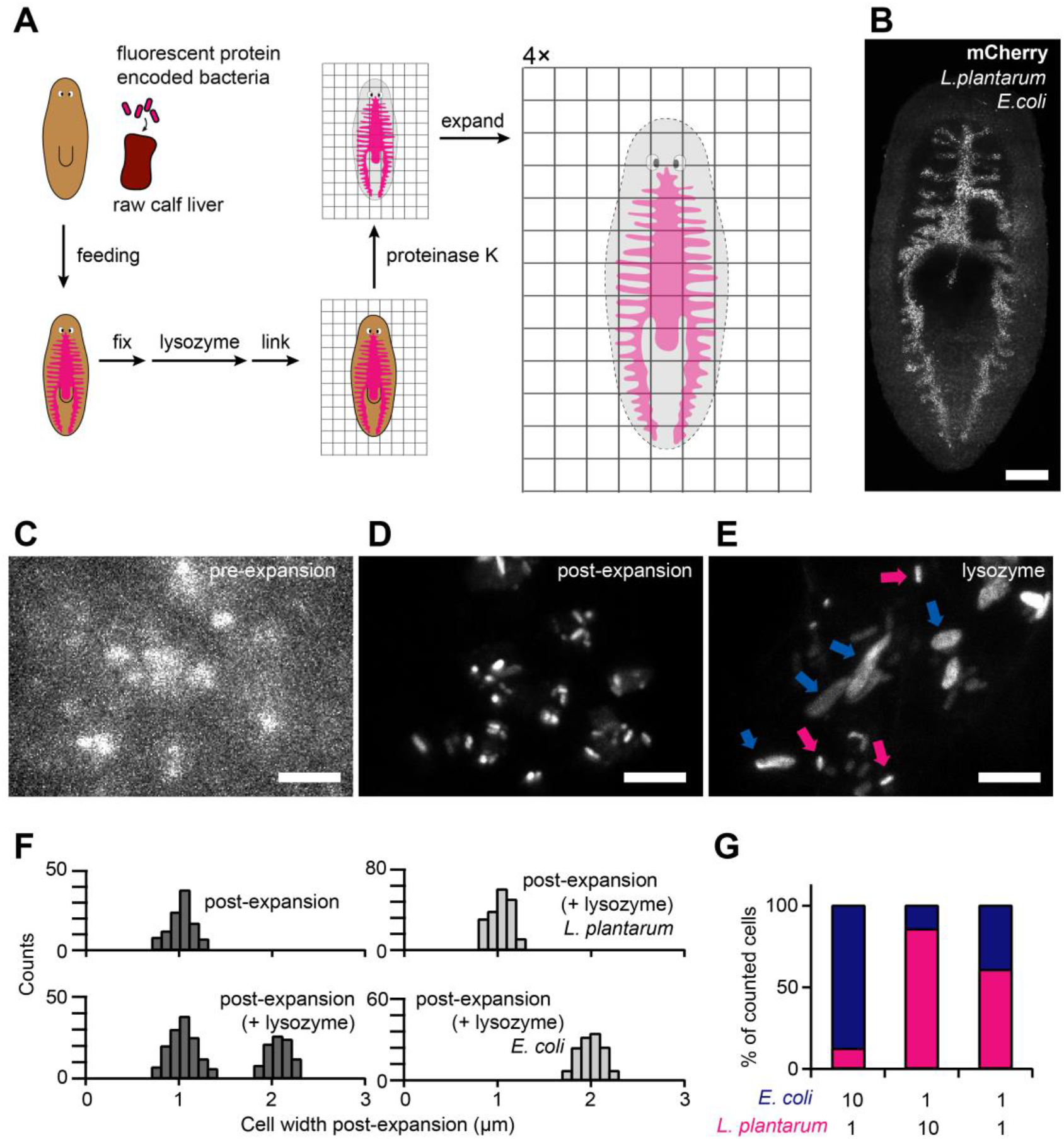
μExM resolves different bacterial species in the planarian flatworm gut. (A) Schematic of the μExM workflow for planarians. Planarians were fed with fluorescent bacteria, and fixed. Unlike other ExM protocols, μExM uses lysozyme or mutanolysin to digest the bacterial cell wall. Linker molecules were then used to anchor the planarian tissue as well as microbial proteins to the hydrogel network. After digestion with proteinase K, the hydrogel was expanded 4-fold isotropically. (B) Pre-expansion maximum-intensity projection of a planarian with its gut colonized by a mixture of *E. coli* and *L. plantarum*, both expressing mCherry. Imaging was performed at 3 d after feeding the planarian with microbes. Scale bar: 200 μm. (C-E) Magnified views showing microbial populations before expansion (C), after expansion (D), and after expansion with lysozyme treatment (E). In (E), magenta arrows indicate unexpanded cells (*L. plantarum*) and blue arrows indicate expanded cells (*E. coli*). Scale bars: 10 μm. (F) Quantification of cell width of the mixed populations of *E. coli* and *L. plantarum* in the planarian gut (left). Right: *In vivo* control populations containing a single species. (G) Species composition in the planarian gut at 3 d post-feeding, counted based on cell width after lysozyme treatment and expansion. *n*>250 cells were measured for each condition. The relative abundance of the two species in the initial mixture fed to the planarians is shown below the plot.

**Figure 4:**
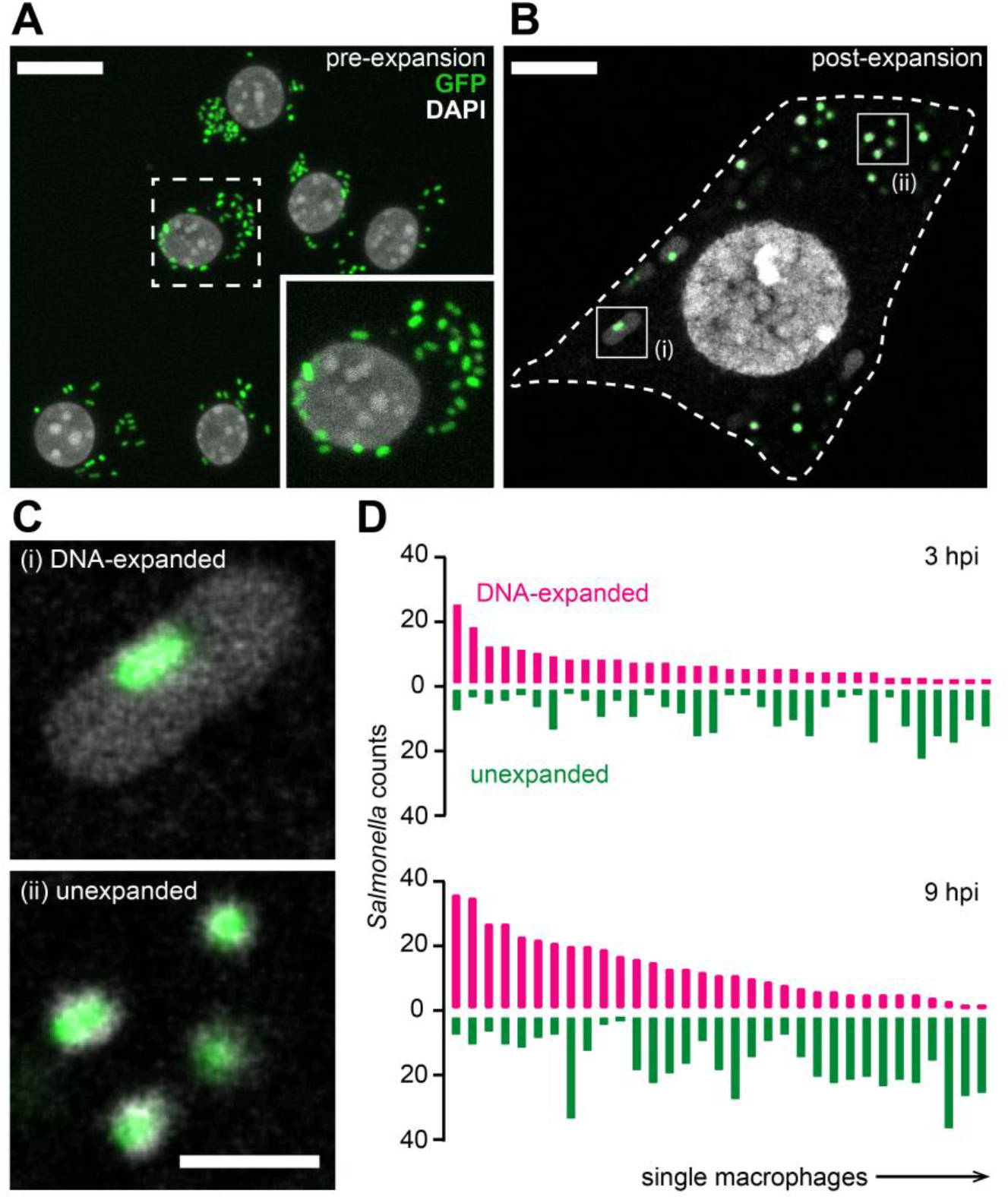
μExM detects changes in the cell-wall structure of macrophage-engulfed *Salmonella* cells. (A,B) Confocal images of RAW264.7 cells infected with GFP-*Salmonella* 3 h post infection before (A) and after (B) expansion. Inset in (A): magnified view of the boxed area. Dashed line in (B): macrophage periphery. Scale bars: 20 μm. (C) Magnified views showing two populations of *Salmonella*: DNA-expanded (top) and unexpanded (bottom), corresponding to cells highlighted by boxes in (B). Scale bar: 5 μm. (D) Number of expanded and unexpanded *Salmonella* cells in individual macrophages determined by manual counting at 3 h (top) and 9 h (bottom) post-infection (hpi). Note that the numbers of both types increase with time as *Salmonella* cells proliferate.

First, we colonized the gut of a model organism, the planarian flatworm *Schmidtea mediterranea* [28], with *E. coli* and *L. plantarum*, both expressing mCherry (**S2 Fig.**). Use of mCherry was prudent as planarian tissues have strong autofluorescence below 560 nm, limiting the utility of other fluorescent proteins such as GFP or YFP that spectrally overlap with the autofluorescence. The bacteria were introduced by feeding the planarian with a calf liver-bacteria mixture, after which colonization was allowed to stabilize for 3 days (**Fig. 3A**). The planarians were then fixed and imaged using the μExM protocol optimized for the planarian tissues (**S3 Fig.**).

Before expansion, bacterial cells in the planarian gut were barely resolvable (**Fig. 3B,C**). Expansion clearly revealed the borders of individual cells, as distances between cells increased (**Fig. 3D** and **S3 Fig.**) and the optical clearing of planarian tissues improved the signal-to-noise ratio (**S3 Fig.**). Moreover, the two species became distinguishable after lysozyme treatment and expansion (**Fig. 3E**); individual cell widths split into two populations corresponding to *E. coli* (2-fold expanded) and *L. plantarum* (approximately unexpanded) (**Fig. 3F**, left). Single-species *in vivo* controls verified that there is little to no overlap between the two populations (**Fig. 3F**, right), with expansion ratios consistent with the *in vitro* measurements (**Fig. 1B**). We quantified the relative abundance of the two species early during colonization, which correlated well with that of the initial mixtures fed to the planarians (**Fig. 3G**). Together, this application demonstrates that μExM can provide quantitative measures of species composition of gut microbiomes, which is critical to resolve the key factors determining these compositions [24,29].

### μExM reveals previously unrecognized cell-to-cell phenotypic heterogeneity among pathogenic bacteria during infection

Next, we investigated *Salmonella* cells during macrophage infection. The fate of *Salmonella* after entering macrophages is known to be heterogeneous: some cells survive and proliferate, whereas others lyse in the harsh intracellular environment [30–32]. Previous studies have suggested that variations in *Salmonella* cell wall structure may play an important role in heterogeneous infection outcomes [33], but it has been challenging to measure such phenotypic variations *in situ*.

We used μExM to image GFP-*Salmonella* cells engulfed by RAW264.7 macrophages, co-staining DNA with DAPI. We observed two types of heterogeneity. First, expansion of individual *Salmonella* cells exhibited two distinct states (**Fig. 4A,B**): some cells remained unexpanded, indicative of an intact cell wall consistent with our *in vitro* experiments (**S4 Figure**), whereas others exhibited an expansion pattern similar to *E. coli imp4213* cells after vancomycin treatment (**Fig. 2**), indicative of nanometer-sized pores in the cell wall through which DNA escaped during expansion to form a halo around the unexpanded cytoplasm (**Fig. 4C**). Second, the fraction of DNA-expanded *Salmonella* cells varied drastically between individual macrophages (**Fig. 4D**). The observed heterogeneity was consistent between time points post-infection. These observations reveal stochasticity in the fate of *Salmonella* cells during macrophage infection, the underlying mechanisms of which will be an important question for future studies.

## Discussion

Here, we develop an expansion microscopy method (μExM) for prokaryotes, and demonstrate that expansion patterns are determined by cell-wall structural properties. We use this phenomenon as a non-conventional imaging contrast, and demonstrate the utility of μExM via *in vivo* imaging of gut microbial communities, and detection of cell-to-cell heterogeneity among pathogenic bacteria as they infect macrophages. We expect this method to spur new research in three major areas.

First, we have shown that in order to be applicable to microbiology, expansion microscopy must be modified to predigest the bacterial cell wall with specific enzymes such as mutanolysin to achieve full uniform expansion. It has been demonstrated that ExM is compatible with conventional antibodies [21,25,34] and RNA fluorescence in situ hybridization [20], therefore μExM is readily adaptable to image nanometer-scale ultrastructures in bacterial cells that are under the diffraction limit of optical imaging. Subcellular organization in bacterial cells is a field of active discovery [35], but has been so far accessible only through specialized super-resolution equipment [22,36,37]. μExM should open up new applications and provide technical convenience in this research area.

More importantly, beyond the established strengths of ExM (i.e., improved spatial resolution and high signal-to-noise ratio), the differential expansion between cells with partially digested cell walls offers a new imaging contrast that is orthogonal to spectral separation in standard fluorescence microscopy using FISH probes, antibodies, or fluorescent proteins. With two relevant applications (*in vivo* gut imaging and macrophage infection), we have demonstrated that the contrast in expansion is quantitative and sensitive to resolve cells of different species or in distinct physiological states, which are otherwise difficult to capture using traditional imaging methods. As μExM requires no special microscopy instrumentation, it can be easily integrated with other optical methods for high-content multimodal imaging. For example, combining expansion and spectral labeling may enable concurrent identification of dozens of microbial species in a complex community or to detect intermediate states as cells undergo physiological changes. We thus anticipate μExM will have broad applications in studies of complex bacterial communities in microbiota, biofilms, and at host-microbe interfaces.

Finally, while cell-wall properties may be measured using direct mechanical methods (e.g., atomic force microcopy [38] or Brillouin microscopy [39]), these methods are not applicable for characterization of cells in dense populations and *in vivo* conditions. μExM can quantitatively evaluate cellular phenotypes *in vivo* and *in situ* under various genetic, chemical, or physical perturbations. These perturbations can include, for instance, genetic disruption of cell-wall synthesis or chemical stresses such as antibiotic treatment [15,40], pH changes, and osmotic shock. Future investigations will be able to compare cellular phenotypes as both isolated individuals and within a dense, complex community. With the recent discovery of micro-scale spatial organization in human microbiota [4,23], μExM will be a powerful tool for revealing how spatial neighborhoods modulate cellular phenotypes.

## Materials and Methods

### Bacterial sample preparation

Bacterial strains and culture conditions used in this study are summarized in **Table 1**. For *in vitro* samples, cells were collected from overnight cultures via centrifugation at 2000 ×*g* for 5 min, washed twice in PBS, and fixed in PBS containing 4% formaldehyde and 1% NP-40 for 10 min. After fixation, cells were washed in PBS and then resuspended in PBST (PBS supplemented with 0.3% Triton X-100) for 30 min at room temperature. The resuspended bacteria were sequentially dehydrated in 50:50% methanol:PBST and then pure methanol.

For vancomycin treatment, *E. coli imp4213* cells were grown to early stationary phase (OD_600_ = 0.8) and incubated in media containing 1 μg mL^−1^ vancomycin (Sigma-Aldrich) for 10 min at 37 °C before fixation. We focused on cells during early stationary phase as we noticed large cell-to-cell variations in expansion between dividing cells during exponential growth.

### Planarian sample preparation

Asexual *S. mediterranea* planarians were maintained at 20 °C in ultrapure water supplemented with 0.5 g L^−1^ Instant Ocean salts and 0.1 g L^−1^ NaHCO_3_ and were fed calf liver paste once or twice weekly. To colonize the planarian gut with bacteria, planarians were starved for at least 7 d, then fed with 250 μL calf liver paste mixed with 50 μL of a mixture of *L. plantarum* and *E. coli* cells, which were collected from 5 mL cultures in early stationary phase (OD_600_ = 0.8–1.0) and concentrated in 1 mL of PBS. After 3 d, individual planarians were collected into separate tubes and fixed individually (to avoid clumping) in PBST containing 4% formaldehyde and 1% NP-40 for 2 h at room temperature. The fixed planarians were washed in PBST, dehydrated in 50:50% methanol:PBST followed by pure methanol, and stored at −20 °C.

To label muscle fibers using immunofluorescence, planarians were killed by 2% HCl for 5 min, and then fixed in 4% formaldehyde with 1% NP-40 for 2 h at room temperature. Samples were rinsed briefly in PBST and bleached overnight at room temperature with 6% H_2_O_2_ in PBST under bright light. The planarians were rinsed with PBST, blocked in PBST supplemented with 1% (w/v) BSA (PBSTB) for 4 h at room temperature, then incubated with the antibody 6G10 (DSHB, 1:1,000 dilutions in PBSTB) for 12–15 h at 4 °C [41]. At least 6 washes of 20 min each with PBST were carried out prior to adding the peroxidase-conjugated secondary anti-mouse antibody (Jackson ImmunoResearch) at a 1:1,000 dilution in PBSTB. After overnight incubation at 4 °C, samples were extensively washed in PBSTB. Tyramide signal amplification was performed by incubating planarians for 10 min in homebrewed TAMRA-conjugated tyramide in 100 mM borate buffer (pH = 8.5) supplemented with 2 M NaCl, 0.003% H_2_O_2_ and 20 μg mL^−1^ 4-iodophenylboronic acid.

### Macrophage sample preparation

To infect macrophages with *Salmonella*, RAW264.7 cells were plated at 500,000 cells/well on coverslips coated with fibronectin (10 μg mL^−1^ for 30 min) in 6-well plates and allowed to attach overnight. Cells were rinsed three times with Fluorobrite DMEM media (ThermoFisher Scientific) supplemented with 10 mM HEPES, 1% FBS, and 2 mM L-glutamine. Overnight cultures of *Salmonella* were diluted in Fluorobrite DMEM media and added to wells at a 1,250:1 multiplicity of infection (MOI). The plate was centrifuged (200 ×*g*) for 15 min at 34 °C. Infected macrophages were washed twice to remove extra bacteria, and 1 mL of medium containing 10 μg mL^−1^ gentamicin was added to each well. Cells were cultured at 37 °C under 5 % CO_2_ for 3-9 h, then fixed in PBS containing 4% formaldehyde and 1% NP-40 for 10 min. After fixation, cells were dehydrated in methanol and stored at −20 °C.

### μExM

Dehydrated samples (bacterial cells, planarian tissues, infected macrophages) were kept at −20 °C for at least overnight and sequentially rehydrated with 50:50% methanol:PBST, then PBST. To digest bacterial cell walls, samples were incubated overnight at 37 °C in either PBS containing 2 mg mL^−1^ lysozyme (ThermoFisher Scientific) or in 50 mM phosphate buffer (pH = 4.9) containing 40 μg mL^−1^ mutanolysin (Sigma-Aldrich), unless otherwise specified.

After cell wall digestion, samples were rinsed three times with PBS, then incubated for 1 h in a PBS solution of 1 mM methacrylic acid *N*-hydroxysuccinimide ester (MA-NHS, Sigma-Aldrich), diluted from a 1 M MA-NHS DMSO stock. After rigorous washes with PBS, samples were incubated in monomer solution (1× PBS, 2 M NaCl, 8.625% (w/w) sodium acrylate, 2.5% (w/w) acrylamide, 0.15% (w/w) N,N’-methylenebisacrylamide) for 1 min (bacterial cells and macrophages) or 45 min (planarian tissues) at 4 °C before gelation.

Gelation was performed in chambers that were assembled using two coverslips as spacers placed between microscope slides. Gelation was initiated by adding monomer solutions supplemented with 0.2% w/w ammonium persulfate stock (10% (w/w), ThermoFisher Scientific) and tetramethylethylenediamine (ThermoFisher Scientific). For planarian tissues and macrophage samples, 4-hydroxy-2,2,6,6-tetramethylpiperidin-1-oxyl (4-hydroxy-TEMPO, Sigma-Aldrich) was added at a concentration of 0.05% (w/w) to inhibit gelation during the diffusion of monomers into tissues. Gelation was completed by incubation at 37 °C for 1 −2 h.

After gelation, gels were gently removed from the chamber and digested overnight at 37 °C in 8 units mL^−1^ Proteinase K (NEB) diluted in digestion buffer (1× TAE buffer, 0.5% Triton X −100, 0.8 M guanidine HCl). Gels were then removed from digestion buffer and placed in excess distilled water to expand. Water was exchanged every 15 min 3-5 times until the size of the expanded gels plateaued. To stain DNA, expanded samples were incubated with 100 μM DAPI or 1 μM TO-PRO-3 for 30 min.

Fluorescence confocal imaging was performed on a Zeiss LSM 800 using either a 20×, N.A. = 1.0, water-immersion objective (W Plan-Apochromat) or a 40×, N.A. = 1.1, water-immersion objective (LD C-Apochromat Corr M27). Expanded samples were mounted in imaging chambers assembled from iSpacer (3.0-mm deep, Sunjin lab). To image a large area, tiled images were stitched using either ZEN or FIJI. The FIJI plugin “Mosaic” was used for the segmentation of bacterial cells. After segmentation, a custom MATLAB script was used to measure aspect ratio and cell width. Cell width is computed along the short axis and averaged at 5 locations evenly spaced along the long axis.

### Estimate of chemical potential for spontaneous translocation of DNA from cell-wall confinement

We model the translocation of a DNA chain through a pore in the cell wall consisting of two primary steps. First, one end of the DNA is anchored near the pore. The anchoring energy and the loss of conformational entropy due to the localization of the chain end gives rise to a free energy barrier of the form [42]

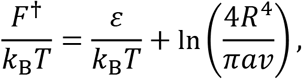

where *ε* is the anchoring energy, *R* the confinement radius (essentially equivalent to the radius of the cell), *a* is the range of anchoring near the pore, and *v* is the volume of the anchored segment. This free energy barrier must be overcome to initialize translocation. Since the anchoring energy depends on the details of the pore and the way it interacts with the anchored segment, it is impractical to evaluate exactly the numerical value of this barrier. Nonetheless, it is apparent that stronger confinement (smaller *R*) lowers the barrier height. Moreover, the presence of multiple pores in the cell wall that are sufficiently large for translocation increases the probability that the DNA is anchored at such a pore.

The second step concerns the actual translocation. After anchored, the DNA chain diffuses along its backbone, outwards or inwards, across the pore. At any instant, one particular segment, labeled *m*, is anchored at the pore, reducing the conformational entropy of the DNA chain (**Fig. 2D**). By treating the chain as a Gaussian random walk, the free energy associated with this entropy loss can be obtained in terms of a series summation [43]. In the limit of strong confinement (i.e., when *R* is smaller than the radius of gyration of the DNA, *R_g_*), the ground-state dominance approximation [42] leads to the following expression for the free energy as a function of *m*:

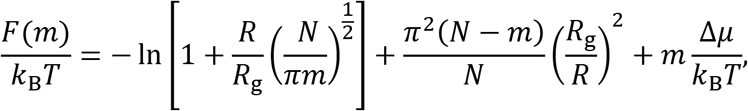

where *N* is the total number of segments and Δ*μ* is the difference in chemical potential of each segment outside and inside the cell. The first and the second terms come from the conformational entropies of the two half-chains outside and inside the cell, respectively. Note that the free energy depends on the confinement through the ratio *R*/*R*_g_.

We evaluated the free energy difference Δ*F*(*m*) ≡ *F*(*m*) – *F*(1) (**Fig. 2E**) for the nucleoid confined inside a bacterial cell. We set the radius of the cell to be *R* = 1 μm, and the length of DNA to be that of *E. coli*. The *E. coli* genome is 4.7 Mbp, having a contour length *L* of 1.596 mm. Using 1 nm as the radius of the cross section, we estimate that the DNA fills ∼0.1% of the intracellular volume, and hence crowding should not substantially hinder segmental motions. Estimating the Kuhn length of DNA at *l*_K_ = 100 nm, we find the number of Kuhn segments *N* = *L*/*l*_K_ = 15,980 and the radius of gyration *R*_g_ = *N*^1/2^*l*_K_ = 5.2 μm. As a result, the DNA is strongly confined, with *R*/*R*_g_ = 0.19.

The salient feature of our model across chemical potential differences Δ*μ* is the existence of a single, *entropic* barrier for molecular translocation (**Fig. 2E**). This barrier originates from the entropy loss associated with translocating the first few segments outside the pore, in addition to those originally anchored inside. Once this entropic barrier is overcome, the free energy decays monotonically with *m*, and DNA translocation proceeds spontaneously. Differentiating the free energy leads to a cubic equation for the location of the barrier, *m*\*, which shows that *m*\* depends on the confinement size *R*/*l*_K_ and on the chemical potential difference Δ*μ*, but *not* on the DNA length *N*. In particular, for Δ*μ* = 0, *m*\* = 0.23(*R*/*l*_K_)^2^; in this case, for *R* = 1 μm, *m*\* = 23, indicating that 23 Kuhn segments (equivalent to ∼6.7 kbp) need to be successfully translocated via diffusive curvilinear motion before the whole chain spontaneously escapes the confinement.

Lowering the segmental chemical potential outside the cell reduces *m*\* and lowers the height of the barrier (**Fig. 2E**). As external localization becomes sufficiently friendly (lower Δ*μ*), the entropic barrier may disappear altogether. By setting *m*\* = 1, we identified that such a transition occurs when

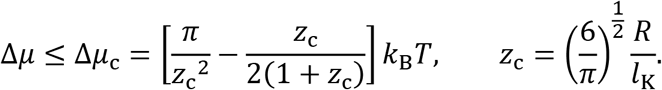

For *R*/*l*_K_ =10, Δ*μ*_c_ = – 0.45 *k*_B_*T* is the chemical potential at which the energy barrier for DNA translocation vanishes.

## Acknowledgements

We thank W. Ludington for sharing the fluorescently labeled *Lactobacillus* and *Acetobacter* strains, with the transformation plasmids originally received under a material transfer agreement from R. Grabherr. We also thank A. Aránda-Díaz, Y. Xue, A. Miguel, M. Rajendram, G. Wong, G. A. O’Toole Jr., K. Y. Han, W. Ludington, and S. Granick for technical help and stimulating discussions. S.B. is supported by a NIH CMB training grant (T32GM007276). This work was supported by the Allen Discovery Center at Stanford on Systems Modeling of Infection (to K.L., K.N., and K.C.H.), a Burroughs Wellcome Fund CASI award (to B.W.), and a Beckman Young Investigator Award (to B.W.). K.C.H. is a Chan Zuckerberg Investigator.

---

**S1 Figure:**
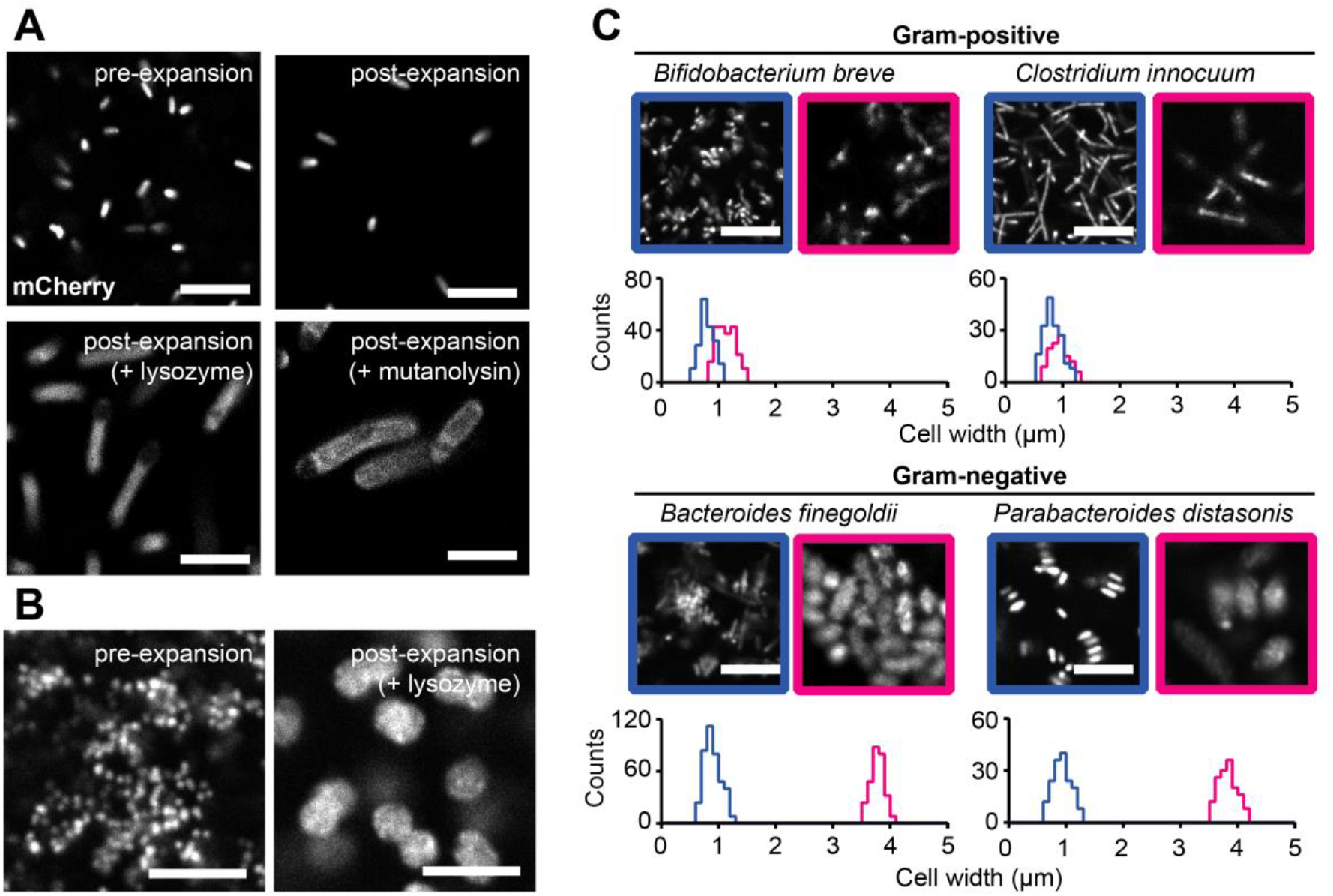
μExM expands microbial species to different extents. (A) Representative μExM images of mCherry-*E. coli*. Corresponding quantifications of cell width with the various treatments are shown in Fig. 1B. (B) μExM images of *Acidaminococcus intestini*, with DNA stained using TO-PRO-3. Corresponding quantifications of cell width are shown in Fig. 1B. (C) Representative μExM images of various human commensal bacterial species. DNA was stained with TO-PRO-3. Blue: pre-expansion images; magenta: post-expansion images after lysozyme treatment. Corresponding quantifications of cell width are shown under the images. All Images are maximum intensity projections. Scale bars: 10 μm.

**S2 Figure:**
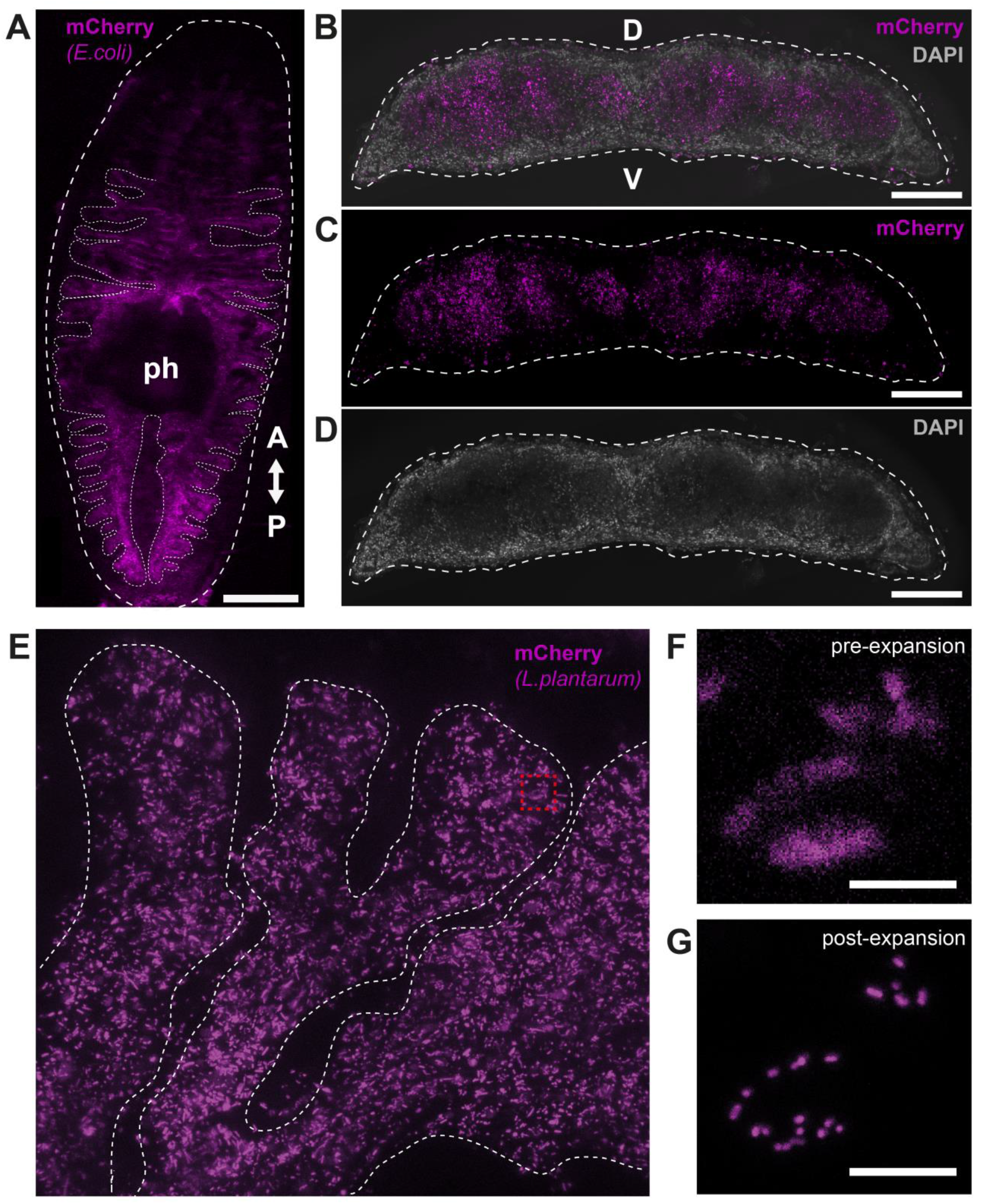
Imaging of fluorescent bacteria colonizing the planarian gut. (A) Confocal image showing a whole planarian, fed with mCherry-*E. coli*, at 3 d post-feeding. The planarian gut and its branches (dotted line) are clearly visible. A, anterior; P, posterior. Scale bar: 500 μm. (B-D) Transverse sections of the planarian trunk region showing that fluorescent *E. coli* are primarily located inside the planarian gut. Dashed line: the outline of the planarian body. D, dorsal; V, ventral. Scale bars: 200 μm. (E) A representative section of the planarian gut colonized b y mCherry-*L. plantarum*, at 3 d post-feeding. Dashed line: the outline of gut branches. (F,G) Magnified views of the highlighted region (red square) in (E), before (F) and after expansion (G). Without cell-wall digestion, *L. plantarum* cells remained unexpanded, but the distances between cells increased 4-fold, allowing single cells to be optically resolved. Scale bars: 10 μm. All images are maximum intensity projections.

**S3 Figure:**
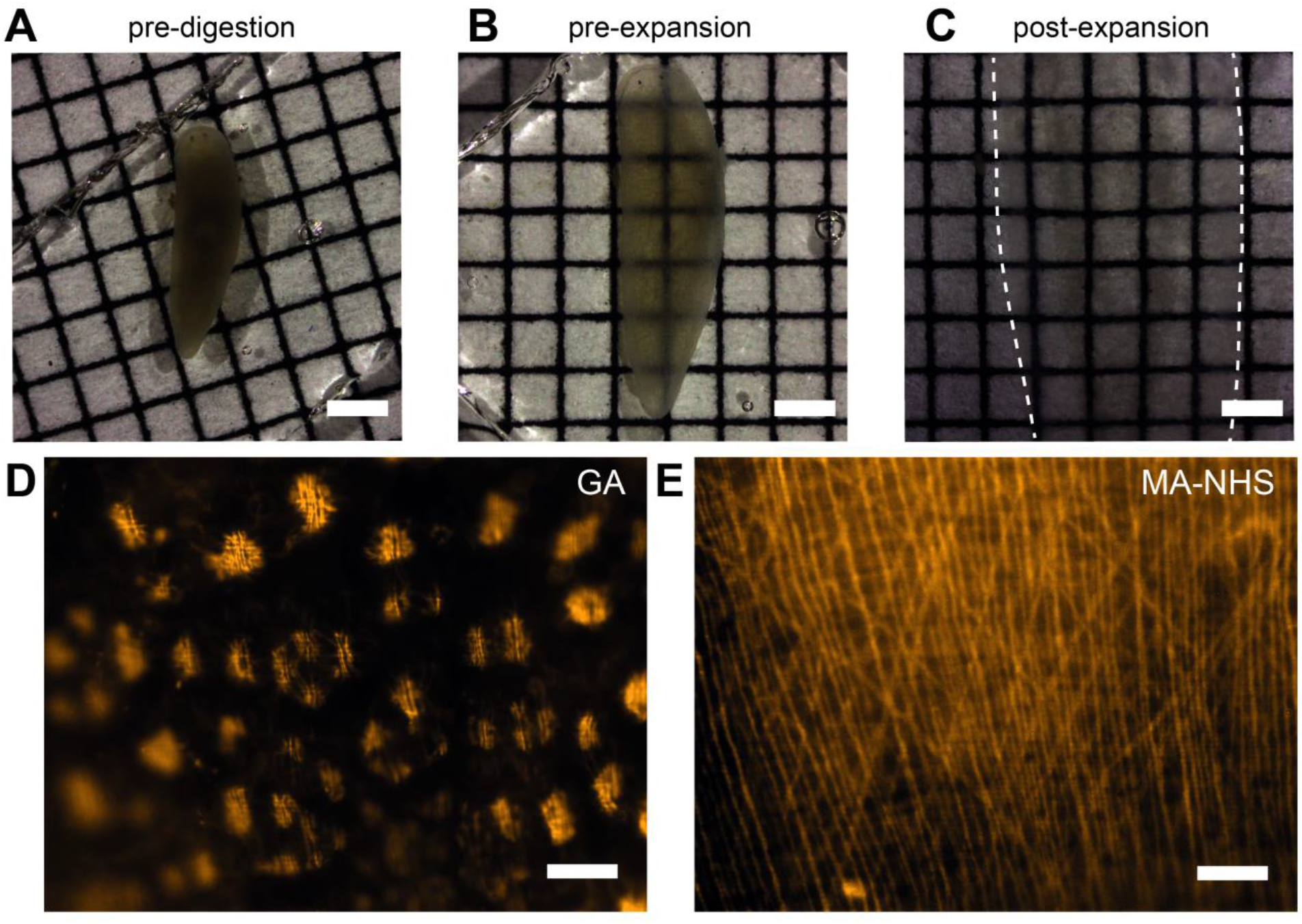
Optimization of μExM for planarian tissues. (A–C) Tissue clearing by digestion and expansion. Grids in the background were included to show tissue transparency. Dashed lines in (C): the outline of the planarian body, which is larger than the imaging view. Scale bars: 1 mm. (D,E) Post-expansion images of planarians immunostained for muscle fibers, using either glutaraldehyde (GA) or MA-NHS as linker molecules. Using GA, expansion disrupts muscle fibers, whereas no distortion was observed in MA-NHS linked tissues. Scale bars: 20 μm.

**S4 Figure:**
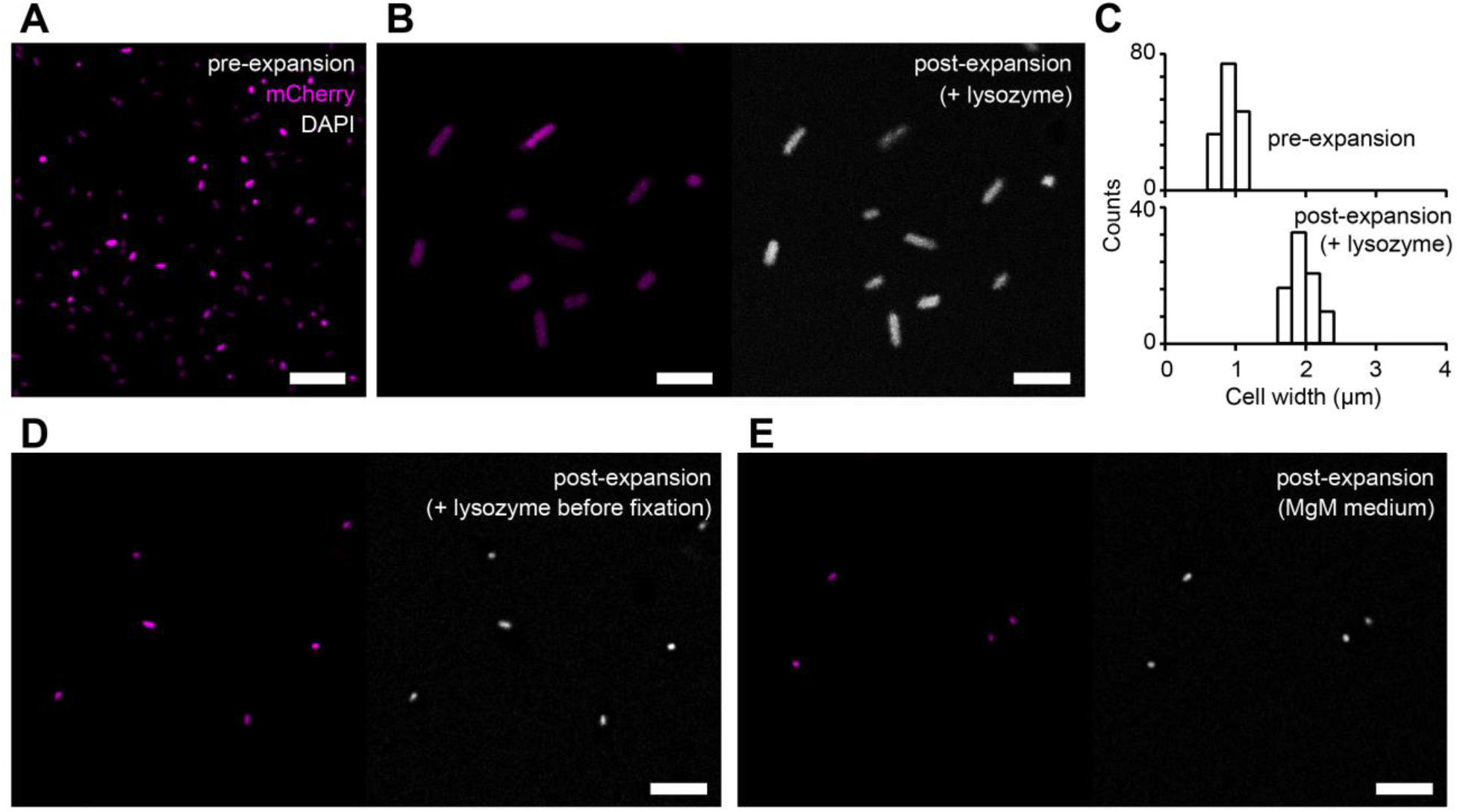
μExM of *Salmonella enterica* cells *in vitro*. (A) Representative maximum intensity projection of mCherry-*Salmonella* cells before expansion. (B) After 1 h lysozyme treatment to digest the cell wall, *Salmonella* cells expanded ∼2-fold. Note that mCherry (left) and DAPI (right) signals co-localized. (C) Quantification of the expansion of cells in images similar to (B). (D,E) Cells treated while alive with 0.5 mg mL^−1^ lysozyme for 1 h at 37 °C prior to fixation (D), or cultured in an acidic, magnesium-depleted minimal medium (MgM-MES, pH 5.0, used to simulate the low pH, low Mg^2+^ environment of the phagosome) (E) did not expand, suggesting that the cell wall remained intact under these conditions. Scale bars: 10 μm.

